# The Histone Demethylase KDM4D Promotes Hepatic Fibrogenesis by Modulating Toll-Like Receptor 4 Signaling Pathway

**DOI:** 10.1101/413245

**Authors:** Fangyuan Dong, Shuheng Jiang, Jun Li, Yahui Wang, Lili Zhu, Xiaona Hu, Yiqin Huang, Xin Jiang, Qi Zhou, Zhigang Zhang, Zhijun Bao

**Author notes:** These authors contributed equally to this work (Fangyuan Dong, Shuheng Jiang). **Correspondence to:** Zhijun Bao, Ph.D., Department of Gastroenterology, Huadong Hospital, Shanghai Medical College, Fudan University, No.221 Yan’an West Road, Shanghai 200040, P.R. China.; or Zhigang Zhang, Ph.D., State Key Laboratory of Oncogenes and Related Genes, Shanghai Cancer Institute, Ren Ji Hospital, School of Medicine, Shanghai Jiao Tong University, 800 Dongchuan Road, Shanghai 200240, P.R. China.; or Qi Zhou, Ph.D., Department of Gastroenterology, Tongji Hospital, Tongji Medical College, Huazhong University of Science and Technology, Wuhan 430030, P.R. China.

## Abstract

Accumulating evidence has revealed the pivotal role of epigenetic regulation in the pathogenesis of liver disease. However, the epigenetic mechanism that accounts for hepatic stellate cells (HSCs) activation in liver fibrosis remains largely unknown. In this study, primary HSCs were used to screen the differentially expressed histone H3 lysine methyltransferases and demethylases during HSC activation. KDM4D was identified as a remarkable up-regulated histone H3 demethylase during HSC activation. The overexpression profile of KDM4D was further confirmed in three fibrosis animal models and human fibrotic liver tissues. *In vitro* genetic silencing of *Kdm4d* impaired the collagen gel contraction and migration capacity of primary HSCs. In established CCl_4_-induced mice model, *Kdm4d* knockdown inhibited fibrosis progression, and promoted fibrosis reversal, with enhanced thinning and splitting of fibrotic septa, as well as a dramatic decrease in collagen area. Whole gene transcriptome analysis showed the regulatory role of KDM4D in Toll-Like Receptor (TLR) signaling pathway. Mechanistically, KDM4D catalyzed histone 3 on lysine 9 (H3K9) di-, and tri-demethylation, which promoted TLR4 expression, and subsequently prompted liver fibrogenesis by activating NF-κB signaling pathways. KDM4D facilitates *TLR4* transcription through demethylation of H3K9, thus activating TLR4/NF-κB signaling pathways in HSCs, contributing to HSC activation and collagen crosslinking, further, hepatic fibrosis progression.

## Introduction

Hepatic fibrosis is caused by various harmful stimuli, characterized by the imbalance of extracellular matrix synthesis, degradation and deposition as well as intrahepatic connective tissue hyperplasia [1-4]. Advanced liver fibrosis eventually leads to irreversible cirrhosis and even hepatocellular carcinoma (HCC), with liver transplantation remaining the only effective treatment for decompensated cirrhosis or advanced HCC [5,6]. Therefore, a better understanding of the molecular mechanisms underlying hepatic fibrogenesis would facilitate the development of preventive and therapeutic approaches for liver fibrosis and possibly for lethal HCC.

The central event during liver fibrogenesis is the activation of hepatic stellate cells (HSCs) [7]. Quiescent HSCs are similar to adipocytes, which can store lipid and retinoid A. Once stimulated, HSCs undergo notable phenotypic transitions, becoming more proliferative and contractile, while *de novo* expression of α-smooth muscle actin (α-SMA) as well as secretion of copious amounts of collagens, which disrupt liver anatomy, herald loss of liver function. This process proceeds to irreversible liver pathology and correlates with augmented mortality of patients with end-stage liver diseases [8,9]. Collagen crosslinking is an essential process for fibrotic matrix stability, which contributes to fibrosis progression and limits reversibility of liver fibrosis. Hence, inhibition of HSC activation and its collagen crosslinking ability are considered to be a promising candidate for halting or even reversing advanced fibrosis.

Epigenetic regulators such as DNA methyltransferases, methyl-DNA binding proteins, histone modifying enzymes and deregulated non-coding RNA have been identified as potential points of therapeutic intervention, which has garnered wide attention [10-12]. Specifically, histone methylation, a reversible process, is one of the most prominent histone posttranslational modifications in response to environmental cues. Accumulating evidence suggests that the methylation of histone lysine residues is a highly dynamic modification owing to the interplay between the epigenetic “writer” lysine methyltransferases (KMTs) and “eraser” lysine demethylases (KDMs) [13-15]. Several KMTs have been reported to be involved in the fibrogenic phenotype of HSC-derived myofibroblasts. For instance, ASH1 orchestrates the coordinated activation of pro-fibrogenic genes including *Acta2, Col1A1*, and *Timp-1* [16]. EZH2 contributes to the transcriptional inhibition of the nuclear receptor PPARγ, which further reprograms the adipogenic HSC towards the myofibroblast phenotype [17,18]. However, the expression patterns and potential regulatory roles of KMTs and KDMs in liver fibrosis require further exploration.

The lysine (K)-specific demethylase 4 (KDM4) family is comprised of 4 isoforms, KDM4A to -D, also known as JMJD2A to -D. KDM4A, B, and C encode proteins consisting of a JmjC, a JmjN, two PHD, and two Tudor domains. KDM4D is unique within the KDM4 family in that it has neither PHD nor Tudor domains, therefore it is only half the size of KDM4A-C [19]. Previous studies showed that KDM4D can be swiftly recruited to DNA damage sites in a PARP1-dependent manner and facilitate double-strand break repair in human cells, which ensure efficient repair of DNA lesions to maintain genome stability [20,21]. KDM4D is also a novel cofactor of androgen receptor since it interacts with androgen receptor and stimulates its ability to up-regulate transcription, which plays an indispensable role in prostate cancer [22].

In this study, a large-scale screen was performed to identify the dysregulated H3 KMTs and KDMs involved during liver fibrosis pathophysiological process. As a result, the most differentially expressed KDM4D, which has never been evaluated in HSC, was selected for detailed investigation. Here we defined the function of KDM4D in HSC activation and liver fibrogenesis.

## Materials and Methods

### Quantitative real-time PCR (qRT-PCR)

RNA extracted from liver tissues and cells was subjected to reverse transcription and subsequently underwent quantitative real-time PCR utilizing 7500 Real-time PCR system (Applied Biosystems, USA). Genes were normalized to β-actin. The relative expression level of each gene was calculated using the formula 2 ^(-ΔΔCt)^. Primer sequences used in qRT-PCR are listed in Supporting Table S1-3.

### RNA interference and gene overexpression

Small interfering RNA (siRNA) oligonucleotides against *KDM4D/Kdm4d, TLR4/Tlr4*, and the scrambled sequences (si-NC) were synthesized by Tuoran Co., LTD (Shanghai, China). The expression vectors pcDNA4-FLAG-KDM4D and its inactive mutant pcDNA4-FLAG-KDM4D-H1122A, which contains histidine-to-alanine point mutation in the JmjC domain and does not possess histone-demethylase function, were synthesized by Sangon Co., LTD (Shanghai, China). For overexpressing TLR4, the pcDNA3.1-TLR4 vector was constructed by GenePharma Co., LTD (Shanghai, China). Constitutively active IKKβ cDNA was synthesized by Sangon Co., LTD (Shanghai, China) and subsequently cloned into pCDNA3.1-Hygro (+) for the experiments. For transfection, LX2 and T6 cells or HSCs were plated at 1 × 10^5^ cells/well in 6-well plates and cultured overnight. Cells were transfected with siRNAs or overexpression vector using RNAimax or Lipofectamine 2000 following the manufacturer’s protocol (Life Technologies, USA). At 72 hours after transfection, cells were harvested for qRT-PCR and western blotting. Primer sequences used for transient interference are listed in Supporting Table S4-7.

### Animal study

Six-week-old male C57BL/6J mice and Sprague Dawley rats were purchased from Shanghai Laboratory Animal Center, Chinese Academy of Sciences (SLAC, CAS). Mice and rats were housed and reared in specific pathogen-free and barrier conditions according to protocols approved by the East China Normal University Care Commission. All animals were housed in a controlled environment under a 12 h dark/light cycle with free access to food and water. All animals received humane care according to the criteria outlined in the “*Guide for the Care and Use of Laboratory Animals*” prepared by the National Academy of Sciences and published by the National Institutes of Health (NIH publication 86-23 revised 1985). All interventions were done during the light cycle.

### CCl_4_, TAA-induced models of non-biliary fibrosis and BDL-induced model of biliary fibrosis

To generate animal models of hepatic fibrosis, 6-week-old male C57BL/6J mice were intraperitoneally (i.p.) injected with carbon tetrachloride (CCl_4_) (dissolved at 1:3 vol/vol in corn oil) (Sigma-Aldrich, St.Louis, MO, USA) or corn oil alone (3.0 ml/kg body weight) twice a week from 6 to 14 weeks of age. Alternatively, 6-week-old male Sprague Dawley rats were i.p. injected with thioacetamide (TAA) (dissolved at 1:3 vol/vol in corn oil) or corn oil alone (0.2 ml/kg body weight) twice a week from 6 to 14 weeks of age. Animals were then divided into test or control groups randomly. Another batch of C57BL/6J mice were subjected to bile duct ligation (BDL) or sham opening operation. They were sacrificed 2 weeks later after operation. All animals were fasted overnight before being sacrificed. Blood samples were harvested for biochemistry analysis. Part of liver tissues was fixed in 10% formalin for paraffin blocks, and the remaining fresh liver tissues were frozen at −80°C for western blotting or qRT-PCR analysis.

### Adeno-associated virus infection

To knockdown *Kdm4d in vivo*, we transduced mice with adeno-associated virus (AAV) serotype 9 that encoded a green fluorescent protein (GFP) reporter together with either short hairpin RNAs targeting *Kdm4d* in liver (sh*Kdm4d*) or empty vector (shNC). The short hairpin RNA (shRNA) sequences targeting the mouse *Kdm4d* gene was cloned into AAV by Genechem Co. LTD (Shanghai, China). Thirty-two C57BL/6J (male, 6-week-old) mice were obtained for this *in vivo* study. After 1 week of acclimatization, they were randomly divided into 4 groups (n = 8 per group) as follows: corn oil+sh-NC, corn oil+sh-*Kdm4d*, CCl_4_+sh-NC, and CCl_4_+sh-*Kdm4d*. The mice were given two i.p. administrations per week of CCl_4_ dissolved in corn oil (1:3 vol/vol) or corn oil alone (3.0 mL/kg) from 6 to 14 weeks of age. At week 12 of age, mice were injected AAV via tail vein (1 × 10^11^ viral particles/mouse) carrying shRNA targeting *Kdm4d* or vehicle alone at one time, followed by additional 2 weeks corn oil or CCl_4_ injection. Mice were sacrificed after overnight fasting, and blood and liver tissues were collected and stored for further analysis.

### Gene expression array

Briefly, LX2 cells transfected with siRNA targeting *KDM4D* or scrambled siRNA were used. Total RNA (15 μg) was isolated from 3 biological replicates and reverse transcribed for hybridization to the whole human Genome Microarray (4 × 44K, Agilent) using an Agilent Gene Expression Hybridization Kit as recommended by the manufacturer. The raw gene expression data were extracted from Agilent Feature Extraction Software (version 10.7) and imported into Agilent GeneSpring GX software (version 11.0) for further analysis. Background subtraction and normalization of probe set intensities were performed using Robust Multiarray Analysis (RMA). Genechip data used for analysis were shown in Supporting Table S8 and S9.

### Protein extraction and western blotting

Western blotting analysis was performed as reported previously [23]. Whole cellular lysates and nuclear fractions were extracted from primary HSCs, activated HSC-T6 and LX2 cells. Nuclear proteins were prepared with the CelLytic NuCLEAR extraction kit (Sigma, NXTRACT, USA) following manufacturer’s introductions. The protein content was determined using a BCA Protein Assay Kit (Beyotime Biotechnology, Shanghai, China). Proteins were subjected to SDS-PAGE and then transferred to a nitrocellulose membrane. After blocking with 5% skim milk for 1 hour at room temperature, the membranes were respectively incubated overnight at 4°C with indicated antibodies shown in Supporting Table S10. After incubation with horseradish peroxidase-conjugated secondary antibodies for 2 hours at room temperature, protein expression was detected by the enhanced chemiluminescent method and imaged with a Bio-Spectrum Gel Imaging System (UVP, USA).

### Chromatin immunoprecipitation (ChIP)

ChIP assay was carried out by using antibodies listed in Supporting Table S11 as well as 100 µg cross-linked native chromatin prepared from primary HSCs and LX2 cells (5 × 10^7^) with or without KDM4D knockdown. Immunoprecipitated DNA was used as the template for quantitative PCR using primers specific for human TLR4 promoter. DNA enrichment was evaluated by average values of the eluate with immunoprecipitated DNA normalized to average values of input.

### Collagen gel contraction assay

Cells at a density of 5 × 10^4^/ml were seeded into 32 mm bacteriological plates (2 ml per dish) in DMEM supplemented with 10% FBS, 1% penicillin and streptomycin and 0.3 mg/ml of acid-extracted collagen I derived from Sprague Dawley rat tail. The cells were cultured at 37°C for 60 min to allow collagen contraction. Then the gels were released from inner edges of plates by tilting plates slightly, and the gel contraction ability was monitored at time points up to 6 h. All assays were performed in triplicate.

### Cell migration assay

Cell migration assays were performed using transwell chambers (Millipore, PIEP12R48, USA). A total of 5 × 10^4^ indicated cells in 200 μl serum-free DMEM were seeded in the upper chamber and 700 μl medium with 10% FBS was added to the lower chamber. After incubation for 24 h, migrated cells were fixed and stained with 0.1% (w/v) crystal violet. Cells were photographed and counted in 5 independent random view fields.

### Statistical analysis

Data are presented as the mean ± standard deviation. Comparisons between two groups were determined by two-sided, unpaired Student’s *t*-test. Comparisons among multiple groups were analyzed by one-way ANOVA test. P values < 0.05 are considered statistically significant.

## Results

### KDM4D expression is up-regulated during HSC activation

Primary HSCs isolated from normal C57BL/6J mice were used to determine the differentially expressed genes of histone H3 KMTs and KDMs during HSC activation. After resting for 24 hours, quiescent (Day 1) HSCs were collected. After culturing for 7 consecutive days, activated (Day 7) HSCs were harvested and analyzed together with Day 1 cells by qRT-PCR. In this study, a total of 19 KMTs and 18 KDMs were tested (Fig. 1A). As a result, the mRNA level of *Lsd1, Ezh1*, and *Kdm6b* were significantly reduced, while the expression level of *Mll1, Mll4, Plu1, Suv39h1, Suv39h2, Riz1, Kdm4a, Kdm4c, Kdm4d, Ezh2, Ash1*, and *Kdm2b* were remarkably increased during HSC activation. Overview of the result, we found that H3K9 KMTs and KDMs were commonly altered. Notably, the mRNA expression level of *Kdm4d* was increased by more than five-fold in activated HSCs when compared with that in the quiescent ones (Fig. 1B). In activated HSCs, *Kdm4d* expression level was much higher than other H3K9 KMTs, including *Suv9h1, Suv39H2* and *Riz1*. From the therapeutic point of view, *Kdm4d* was selected as a candidate gene for further investigation. By co-staining of KDM4D and α-SMA in primary HSCs derived from C57BL/6J mice, we further confirmed that activated HSCs gave rise to KDM4D nuclear expression (Fig. 1C). The qRT-PCR and western blotting also illustrated that KDM4D expression level augmented with HSC activation in a step-wise manner (Fig. 1D, 1E). Together, these data strongly suggested that KDM4D overexpression occurs gradually during HSC activation.

**Figure 1.**
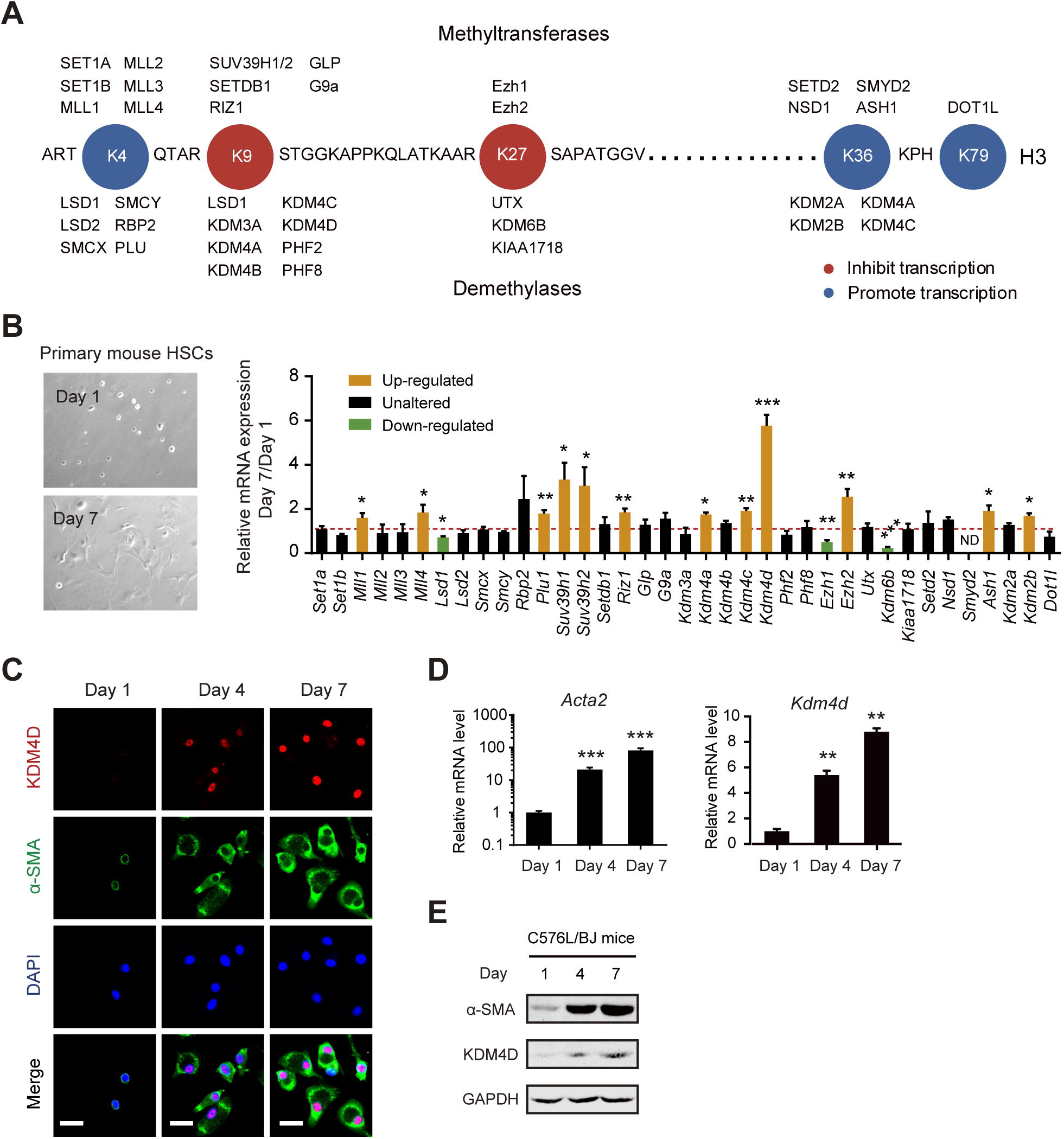
KDM4D expression is up-regulated during HSC activation. (A) Schematic of human histone H3 methyltransferases and demethylases. (B) Representative bright field images of C57BL/6J mouse primary HSCs on D1 and D7 (left panel). Relative mRNA expression levels of H3 methyltransferases and demethylases in primary HSCs (D7/D1) of C57BL/6J mouse (right panel). ND, not determined. (C) Cellular immunofluorescence staining of KDM4D and α-SMA in primary HSCs of C57BL/6J mice on day 1, 4, and 7. Scale bar, 25 μm. (D, E) The mRNA and protein expression level of *Kdm4d* and *Acta2* in primary HSCs of the C57BL/6J mice on day 1, 4, and 7. Data were presented as mean ± SD, *p < 0.05, **p < 0.01, ***p <0.001.

### KDM4D expression is up-regulated during biliary and non-biliary fibrogenesis

To further validate the expression pattern of KDM4D in liver fibrogenesis, biliary and non-biliary murine models induced by CCl_4_ (left), TAA (middle), and BDL (right) were generated (Fig. 2A). Repeated CCl_4_ and TAA administration led to robust and progressive scarring, characterized histologically as significantly pericentral fibrosis, while BDL contributed to biliary fibrosis. As shown in Fig. 2A, compared with their control liver sections, the fibrotic liver tissues had larger fibrotic areas (H&E and α-SMA staining), and severer collagen deposition (Sirius Red and Masson staining), respectively. As expected, IHC revealed that KDM4D, which was hardly detectable in non-fibrotic healthy livers, was strongly expressed after BDL operation or consecutive CCl_4_/TAA injection (Fig. 2A). The intense immunoreactivity of KDM4D in the fibrotic areas was widely observed in all the three animal models indicate of a common expression profile of KDM4D in both parenchymal and biliary fibrosis. Furthermore, RNA was isolated from primary HSCs from CCl_4_/TAA/oil or BDL/sham animal livers and we found that *Kdm4d* was commonly and highly expressed during liver fibrogenesis (Fig. 2B-D). To estimate the clinical relevance, we then performed dual immunofluorescence staining of KDM4D expression in human fibrotic liver tissue samples. Interestingly, KDM4D is was specifically expressed in the fibrous septa and mainly co-localized with α-SMA in fibrotic liver tissues, implying the restricted expression of KDM4D in myofibroblast-like cells in liver fibrosis (Fig. 2E).

**Figure 2.**
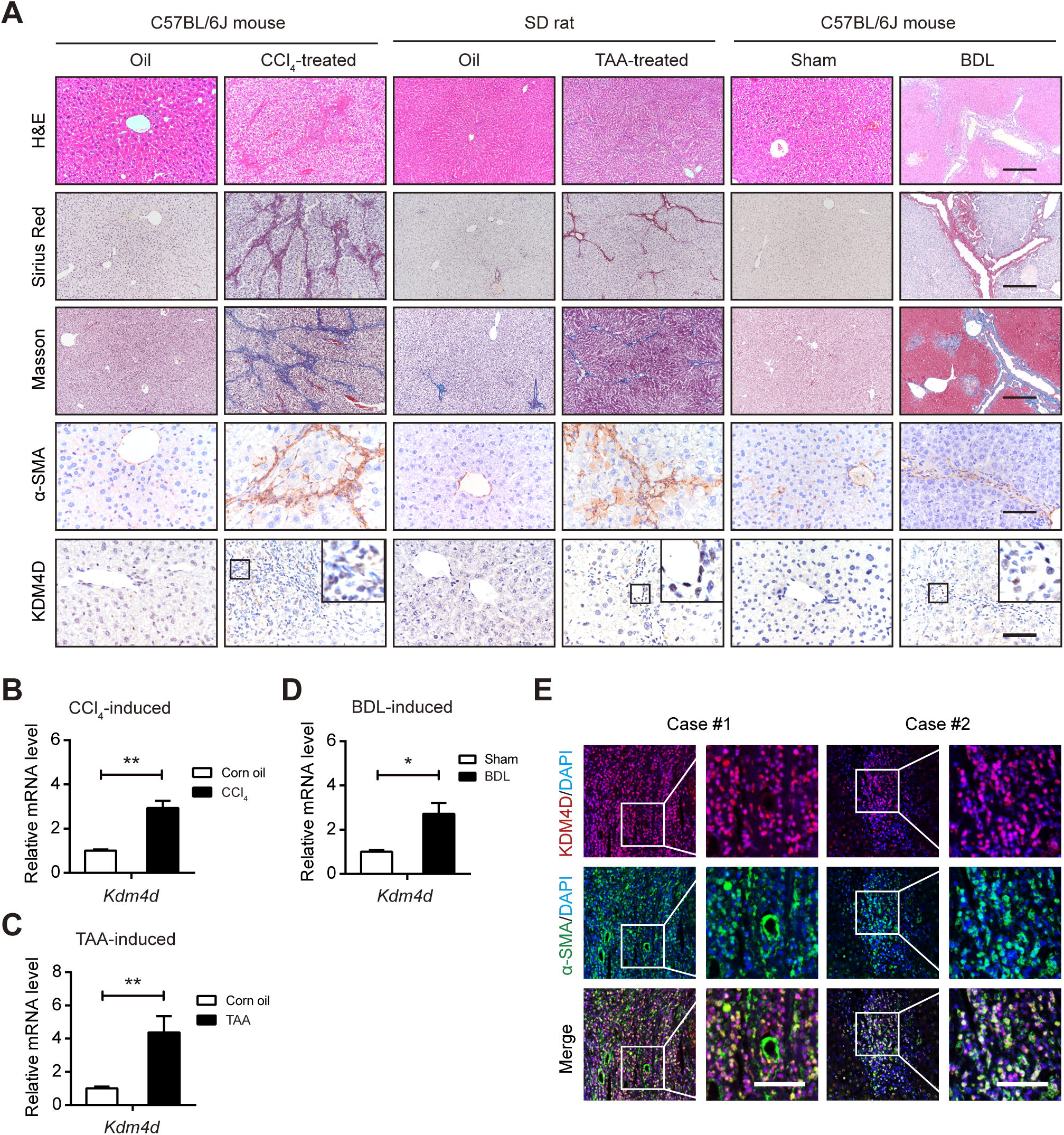
KDM4D is up-regulated during biliary and non-biliary fibrogenesis. (A) Representative images of H&E, Sirius red, Masson’s trichrome staining, α-SMA, and KDM4D staining in liver tissues from CCl_4_-induced C57BL/6J mice, TAA-induced SD rat, and BDL-induced C57BL/6J mice as well as their corresponding counterparts. Scale bar, 200 μm for H&E, Sirius red, and Masson’s trichrome staining; 100 μm for α-SMA and KDM4D staining. (B-D) Real-time qPCR analysis of *Kdm4d* mRNA level in quiescent or activated HSCs isolated from CCl_4_/TAA/BDL/Control animal livers. (E) Representative immunofluorescence con-focal images of α-SMA (green), KDM4D (red), and DAPI (blue) in tissue sections from human fibrotic liver. Scale bar, 50 μm. Data were presented as mean ± SD, *p < 0.05, and **p < 0.01.

### *Kdm4d* deficiency ameliorates liver fibrosis in mice

Next, we were prompted to investigate the *in vivo* role of *Kdm4d* knockdown in a CCl_4_-induced liver fibrosis mouse model (Fig. 3A). *Kdm4d* knockdown in liver tissue was then carried out using tail vein injection with a *Kdm4d* shRNA-expressing AAV9. AAV9 targets many tissues including central nervous system, heart, liver, and lung. We therefore tested the specificity of KDM4D knockdown in liver tissues. To address this, resident liver cells, including hepatocytes, sinusoidal endothelial cells (SEC), Kupffer cells (KC), portal myofibroblasts (PF), and HSCs from CCl_4_/oil mouse livers were isolated. RT-qPCR result showed that *Kdm4d* expression was mainly derived from activated HSCs, indicating the specificity of AAV9-mediated *Kdm4d* knockdown in HSCs (Fig. 3B). Indeed, AAV9-mediated *Kdm4d* deficiency led to significant reduction of KDM4D protein in fibrotic liver tissues (Fig. 3C). *Kdm4d* deficiency had no obvious deleterious effect on normal liver function, but significantly alleviated CCl_4_-induced liver injury (Fig. 3D). Specifically, *Kdm4d* knockdown exhibited a reduction in collagen accumulation, compared with their counterparts in the control group (Fig. 3E, F). Consistently, lower expression of α-SMA was found in *Kdm4d*-deficient mice than their control littermates as revealed by IHC analysis (Fig. 3G). Collectively, these *in vivo* data reinforced the notion that KDM4D served as an important driving force underlying the pathogenesis of liver fibrosis.

**Figure 3.**
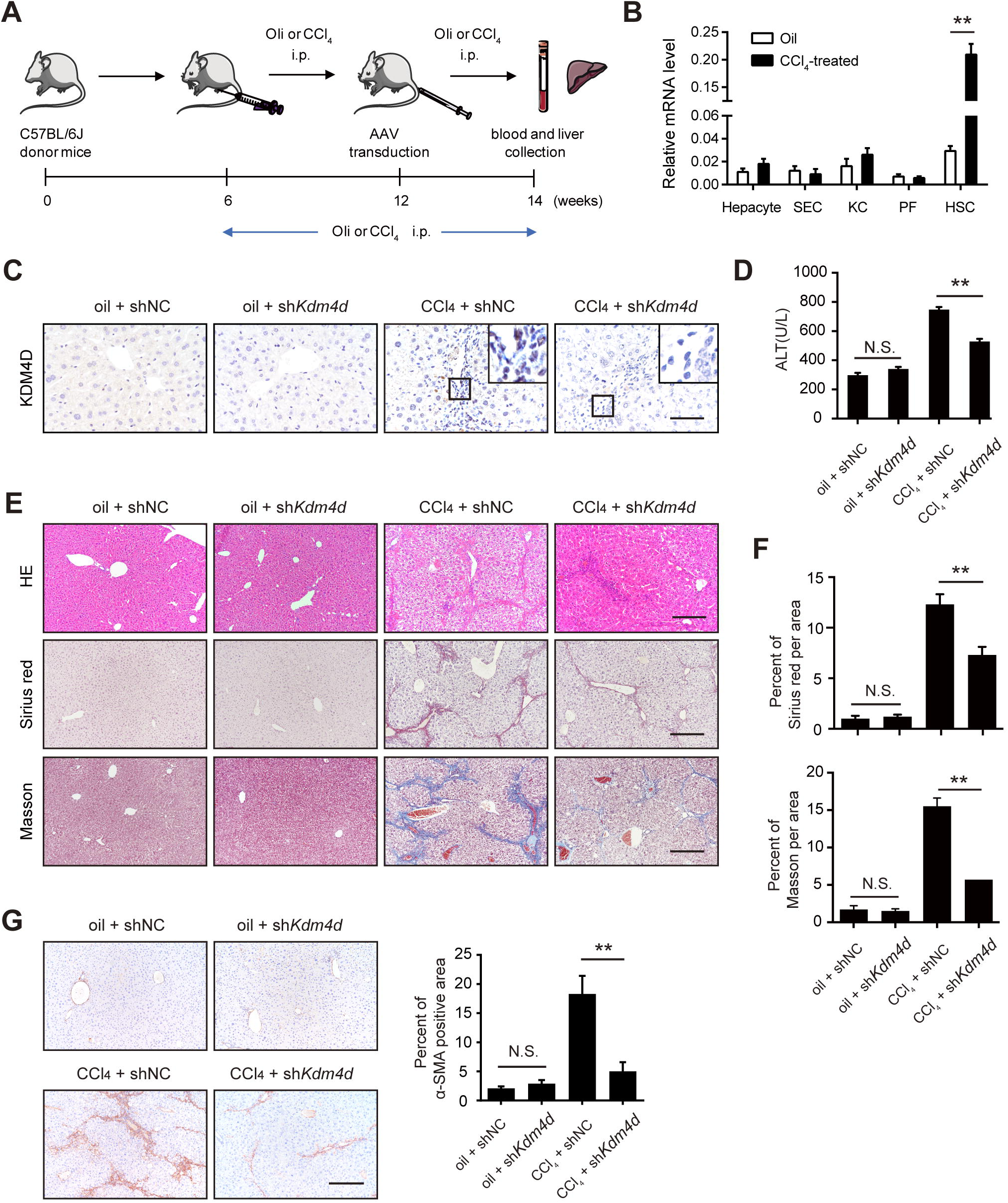
*Kdm4d* deficiency ameliorates liver fibrosis in mice. (A) The experiment scheme for *Kdm4d* knockdown *in vivo*. Liver fibrogenesis was induced by CCl_4_ injury using intraperitoneal injection twice a week from 6 to 14 weeks of age. Their counterparts were injected with oil. At week 12, shRNA expression vectors GFP carrying shRNA targeting *Kdm4d* or negative control were transfected into mouse liver through tail vein injection. Animals were humanely sacrificed, and their blood and liver samples were collected at week 14. (B) RT-qPCR analysis of *Kdm4d* mRNA level in HSCs, hepatocytes, sinusoidal endothelial cells (SEC), Kupffer cells (KC), and portal myofibroblasts (PF) isolated from the normal or CCl_4_-indced mouse liver tissues. (C) Representative IHC images of KDM4D in indicated groups. Scale bar, 200 μm. (D) The liver injury in each group was measured by ALT levels. (E) Representative histology images of H&E, Sirius red and Masson’s trichrome staining in indicated groups. Scale bar, 200 μm. (F) Sirius red and Masson’s trichrome positive areas were measured by Image J software. (G) Representative IHC images of α-SMA in indicated groups. Positive α-SMA staining area was measured by Image J software. Scale bar, 200 μm. Abbreviation: NS: not significant, shNC, empty vehicle alone. Data were presented as mean ± SD, **p < 0.01.

### *Kdm4d* knockdown effectively suppresses HSC activation

As HSCs are of vital importance in liver fibrogenesis, we further probed the impact of *Kdm4d* deficiency on HSC activation *in intro*. HSCs from sh*Kdm4d* mice had a pronounced depletion of KDM4D compared with that in the control mice as detected by western blotting (Fig. 4A). Likewise, transcriptional silencing of *Kdm4d* also dramatically diminished the expression of the pro-fibrogenic genes in primary HSCs including *Acta2, Col1a1*, and *Vim* as monitored by qRT-PCR (Fig. 4B). Moreover, immunofluorescence staining displayed that KDM4D knockdown attenuated the fibrogenic phenotype of activated HSCs (Fig. 4C). Given that activated HSCs are characterized by enhanced migratory and contractile capacities, we further investigated the impact of *Kdm4d* knockdown on these properties. Indeed, *Kdm4d* knockdown significantly impaired the collagen gel contraction and migratory capacities of primary HSCs as demonstrated by collagen gel contraction assay (Fig. 4D) and transwell cell migration assay (Fig. 4E), respectively. Meanwhile, loss-of-function studies in LX2 and T6 cells confirmed similar observations (Supporting Fig. S1). These observations clearly demonstrate that KDM4D might be essential for HSC activation and *Kdm4d* deficiency could attenuate the crosslinking function and migration capacity of HSCs in mice.

**Figure 4.**
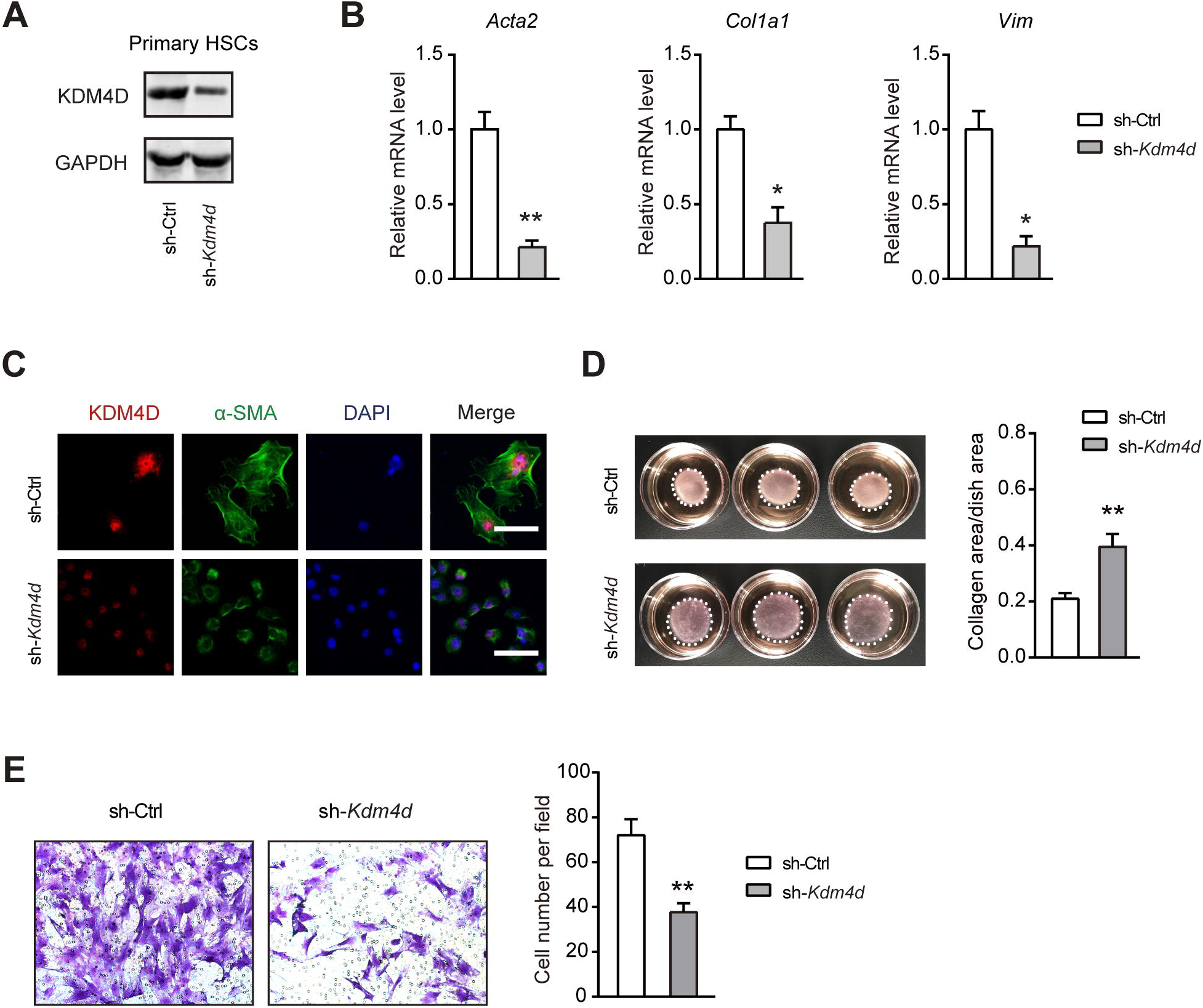
*Kdm4d* knockdown effectively suppresses HSC activation *in vitro*. (A) Western blotting analysis of *Kdm4d* knockdown efficiency in activated C57BL/6J mice primary HSCs. (B) Effects of *Kdm4d* knockdown on the expression of pro-fibrogenic genes (*Acta2, Col1a1*, and *Vim*) were examined by qRT-PCR. (C) Representative immunofluorescence images of primary HSCs from sh-Ctrl and sh-*Kdm4d* mice. (D) Collagen gel contraction assay of primary HSCs from sh-Ctrl and sh-*Kdm4d* mice (n = 3 for each group). Statistical analysis of collagen area/dish area was shown in the right panel. (E) Representative migration images of primary HSCs from sh-Ctrl and sh-*Kdm4d* mice (n = 5 for each group). Statistical analysis of cell number per field was shown in the right panel. Data were presented as mean ± SD, *p < 0.05, and **p < 0.01.

### KDM4D modulates TLR4 expression in HSCs

To uncover the mechanism by which KDM4D promotes liver fibrosis, a whole genome transcriptomic analysis was performed in si-NC or si-*KDM4D* treated LX2 cells. ClueGO analysis of the differentially downregulated genes (si-*KDM4D* vs. si-NC) revealed the predicted biological associations of these altered genes including the Toll-like receptor (TLR) signaling pathway, intestinal immune network for IgA production, rheumatoid arthritis, transcriptional deregulation in cancer, African trypanosomiasis, leukocyte transendothelial migration, and cytokine-cytokine interaction (Fig. 5A and Supporting Fig. S2A). It is well known that TLR4 signaling is critically involved in the activation of HSCs during liver injury. Indeed, knockdown of *TLR4* in LX2 cells dramatically inhibited the mRNA expression of pro-fibrogenic genes (Supporting Fig. S2B, S2C). We therefore hypothesized that KDM4D might activate TLR4 expression by epigenetic modifications (Fig. 5B). Indeed, *Kdm4d* knockdown mainly led to dramatic reduction of TLR4 protein level and enhanced remarks of H3K9me2 and H3K9me3, but not H3K9me1 (Fig. 5C). The ChIP assay validated that compared with the negative control (mouse IgG), marked enrichment of the *TLR4* promoter was found in the cistrome of H3K9me2 and H3K9me3 in *KDM4D* knockdown LX2 cells (Fig. 5D). Consistently, H3K9me2 ChIP analysis confirmed much higher H3K9me2 markers on the *TLR4* promoter in KDM4D knockdown cells (Fig. 5D). Moreover, KDM4D ChIP analysis showed that *KDM4D/Kdm4d* knockdown markedly attenuated the occupancy of KDM4D on the *TLR4/Tlr4* promoter in the LX2 cells (Fig. 5E) and primary HSCs (Fig. 5F), respectively. We also analyzed the implication of *KDM4D* overexpression on *TLR4* expression by transient transfection of LX2 cells with extraneous expression vectors for *KDM4D* and its demethylation-defective mutant. As a result, overexpression of *KDM4D* but not the demethylation-defective mutant form significantly up-regulated the mRNA and protein expression of *TLR4* (Fig. 5G, 5H). These observations clearly demonstrate the epigenetic regulatory role of KDM4D on the expression of TLR4 through modulating H3K9me2 and H3K9me3 status on its promoter in HSCs.

**Figure 5.**
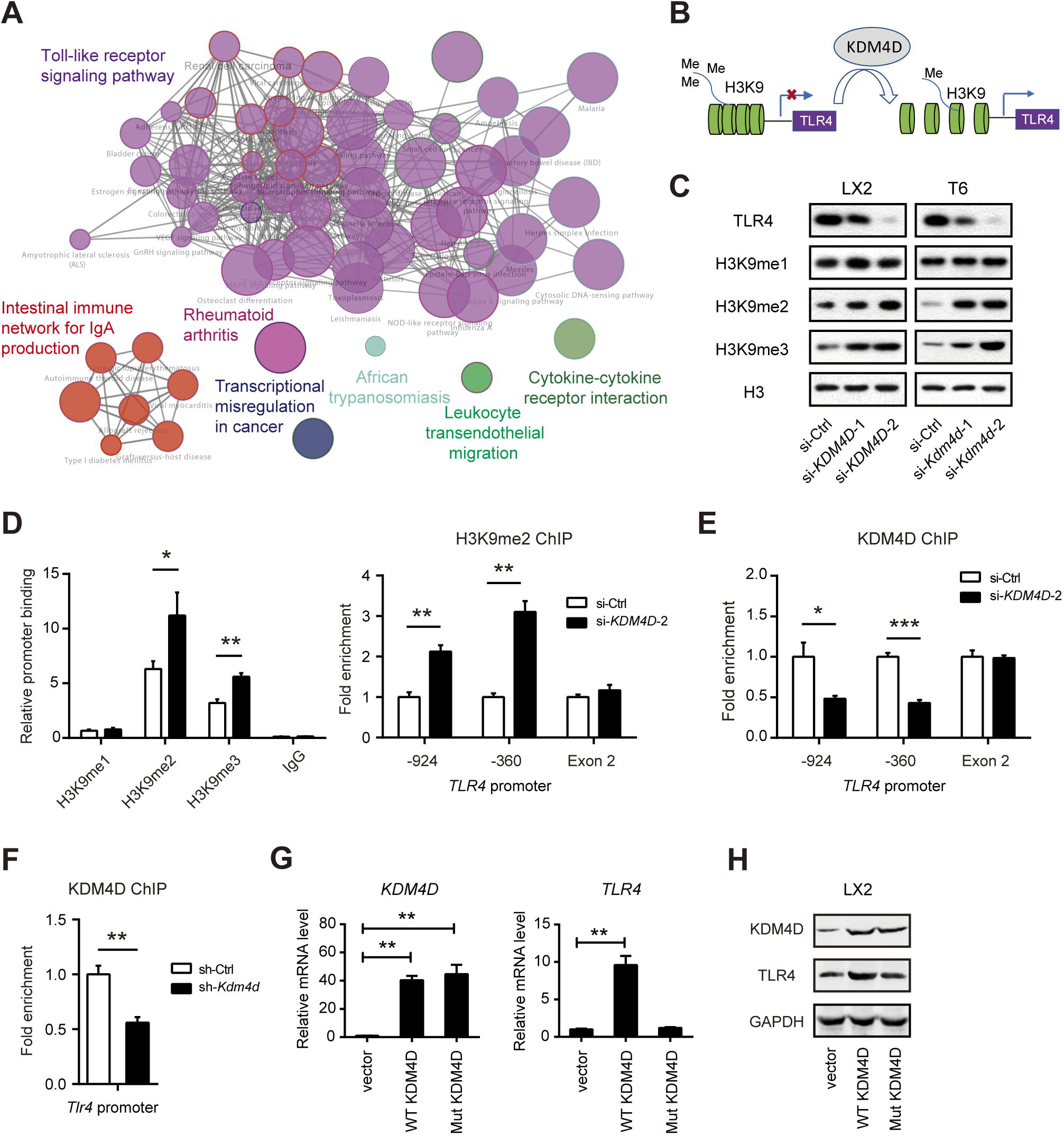
KDM4D modulates *TLR4* expression in HSCs. (A) Expression profile chip analysis with scramble siRNA (NC) or si-*KDM4D*; ClueGO analysis of the differentially downregulated genes by CytoScape software. Enriched pathways are shown as nodes interconnected based on the κ score. (B) Model depicting the role of KDM4D in transcriptional regulation of *TLR4* expression in liver fibrogenesis. (C) Immunoblotting of TLR4, H3K9me1, H3K9me2, H3K9me3 expression in LX2 and T6 cell lines transfected with specific siRNAs against *KDM4D/Kdm4d*. Histone H3 serves as the loading control. (D) ChIP assays were performed with indicated antibodies in LX2 cells with or without the *KDM4D* knockdown, and enrichment of target DNA was analyzed with qPCR using primers specific for the *TLR4* promoter. Fold enrichments were calculated by DNA enrichment bound with H3K9me2 antibody compared with those bound with IgG. (E) KDM4D ChIP assays were performed in LX2 cells with or without *KDM4D* knockdown, and enrichment of target DNA was analyzed with qPCR using primers specific for the *TLR4* promoter. (F) KDM4D ChIP assays were performed in primary HSCs isolated from mice with or without AAV-sh*Kdm4d* treatment, and enrichment of target DNA was analyzed with qPCR using primers specific for the *Tlr4* promoter. (G, H) Effects of overexpressing of KDM4D and its demethylation-defective mutant vector on the mRNA (G) and protein expression (H) of *TLR4* in LX2 cells. Data are presented as mean ± S.D. *p < 0.05, **p < 0.01, and ***p < 0.001.

### KDM4D promotes liver fibrosis by activating TLR4/NF-κB signaling pathways

Upon stimulation with ligands, such as Lipopolysaccharides (LPS), TLR4 receptor triggers the myeloid differentiation factor 88 (MyD88)-dependent pathway and activates downstream nuclear factor κB (NF-κB) pathways (Fig. 6A) [24]. NF-κB is the master regulator for the transcription of pro-inflammatory mediators in the liver [25]. Consistently, the protein expression of MyD88 and phosphorylated p65 were significantly increased in isolated HSCs of CCl_4_-treated group, but notably declined when *Kdm4d* was knocked down *in vivo* (Fig. 6B). Additionally, KDM4D deficiency also significantly ameliorated the protein expression of α-SMA and TLR4 in isolated HSCs upon LPS stimulation as revealed by western blotting (Fig. 6B), suggesting the regulatory role of KDM4D in liver fibrosis might be mediated by TLR4-NF-κB signaling pathway. To confirm this hypothesis, we silenced *Kdm4d* and overexpressed *Tlr4* in the primary isolated HSCs (Fig. 6C). Excitingly, we noticed that KDM4D deficiency in activated HSCs significantly decreased NF-κB transcriptional activity, which can be elevated by overexpression of *Tlr4* (Fig. 6D). Rescue experiments also showed that decreased expression of pro-fibrotic genes (Fig. 6E), impaired contractile capacity (Fig. 6F), and migratory ability (Fig. 6G) of primary HSCs induced by *Kdm4d* knockdown can be largely restored by introduction of *Tlr4*. Likewise, similar observations were also found in LX2 cells (Supporting Fig. S3). Together, these observations demonstrate that that KDM4D contributes to liver fibrosis possibly through modulating TLR4/NF-κB signaling pathways.

**Figure 6.**
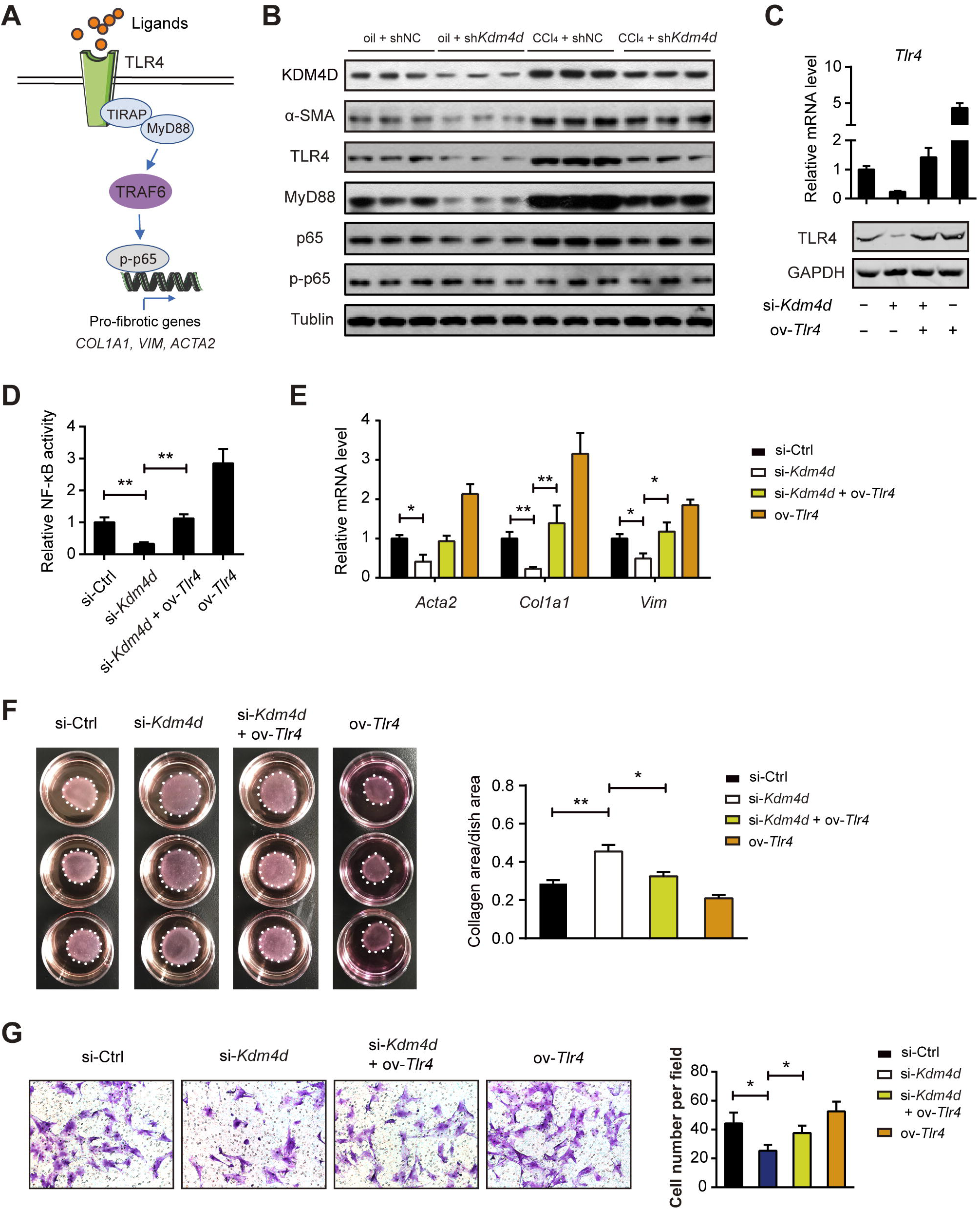
KDM4D promotes liver fibrosis by activating TLR4/NF-κB signaling pathways. (A) Schematic depicting TLR4/MyD88/NF-κB signaling pathway. (B) Proteins from primary HSCs in oil + shNC, oil + sh*Kdm4d*, CCl_4_ + shNC, and CCl_4_ + sh*Kdm4d* groups were harvested and assayed by western blotting (n = 3 representative mice for each group) with indicated antibodies. Tubulin was used as the loading control. (C) Certification of *Kdm4d* and *Tlr4* expression at both mRNA and protein level in the presence of *Kdm4d* knockdown and/or *Tlr4* overexpression in primary isolated HSCs. (D-G) The effects of *Tlr4* overexpression on the NF-κB transcriptional activity, expression of pro-fibrotic genes, and contractile and migratory capacities in *Kdm4d* knockdown primary HSCs. Data are presented as mean ± S.D. *p < 0.05, **p < 0.01.

## Discussion

Hepatic fibrosis, characterized by HSC activation and excessive production and deposition of extracellular matrix in the liver, serves to disrupt normal liver structure and heralds more severe, irreparable liver pathologies such as hepatic failure and hepatocellular carcinoma. HSCs, the major source of fibrogenesis in the liver, locate in the Disse pace of the normal liver, and are activated into myofibroblasts by chronic stimulation. Once activated, HSCs show enhanced cell proliferation, express excessive α-SMA, and overproduce extracellular matrix. Post-translational regulations in mammals are dictated by the epigenetic mechanism, of which histone modifications constitute a key branch. They can influence chromatin structure, ultimately leading to gene expression alterations [26]. A series of epigenetic enzymes are actively involved in the addition or removal of covalent modifications, which include acetylation/deacetylation, methylation/demethylation, phosphorylation, ubiquitination, and sumoylation [27,28]. Deregulation of these processes is a hallmark of pathogenesis. The mechanism of epigenetic regulation in hepatic fibrogenesis has not been well elucidated, therefore, studies on the relationship between epigenetic modifications and liver fibrosis will refine our understanding of the fibrotic pathogenesis. The complexity of dynamic regulatory networks in the pathogenesis of liver fibrosis raises the importance of further exploration for novel factors and signaling pathways. Here, we uncovered that KDM4D, a histone demethylase, as a novel epigenetic regulator of HSC activation and thereby modulates hepatic fibrogenesis by altering methylation status of H3K9.

KDMs are versatile proteins that modulate multiple cellular processes, such as gene expression regulation, cell differentiation, embryonic stem cell renewal, and tumor development [29]. Our data portray KDM4D as a protein bridging H3K9 demethylation with HSC activation. Using H3K9me2 as a proxy for KDM4D activity is reasonable, as its level is coregulated by both histone methyltransferases and demethylases. Coincidently, the level of both histone H3 methyltransferases and demethylases are upregulated during HSC activation. Given KDM4D is the most abundant histone demethylase whose expression level is much higher than any of the other histone methylase or demethylase, we can say that it is histone demethylases rather than methyltransferases who predominantly contribute to the decline of H3K9me2 and H3K9me3 level during HSC activation. KDM4D consists of evolutionarily conserved JmjN and JmjC domains at its N terminus whereas the overall sequence of its C-terminal region contains no obvious characterized domain [10]. Apart from its well-known regulatory roles in the DNA damage response, KDM4D also stimulates p53-dependent transcriptional activity, which points to a pro-oncogenic implication [30]. In this study, a novel function of KDM4D in liver fibrogenesis was described. We uncovered that the expression of KDM4D greatly increased during trans-differentiation of quiescent HSCs to activated ones, also it was required for collagen contraction and migration capacity of HSCs. Further, the severity of established liver fibrosis was alleviated as a result of *Kdm4d* knockdown *in vivo. Kdm4d*-deficient mice showed a notable reduction in not only hepatic injury severity. At the same time, *Kdm4d* depletion caused a significant suppression of crosslinked collagen deposition, which derived from HSC activation.

TLRs were originally identified as pathogen-associated molecular pattern recognition receptors that recognized exogenous ligands in response to infection [31]. In cirrhotic mice or patients, the gastrointestinal tract produces and absorbs considerable bacterial LPS with increased permeability of the intestinal mucosal barrier. It is well known that TLR4 signaling plays a pivotal role in liver inflammation and fibrosis. Therefore, targeting TLR4 signaling pathways represents an attractive strategy for liver fibrosis treatment [32]. In this study, our data portrayed that differentially expressed KDM4D modulated TLR4 level in HSCs by modulating chromatin structure. We found that genetic silencing of *Kdm4d* in HSCs resulted in a significant inhibition of TLR4 signaling pathway and KDM4D promoted *TLR4* transcription through its demethylation activity. Engagement of ligands with the TLR4 receptor induces activation of intracellular signaling pathways through recruitment of the receptor adaptors MyD88, resulting in activation of the IκBα kinase complex and subsequent translocation of NF-κB (p65) [24]. Consistently, the expression level of TLR4 and the phosphorylated p65 as well as the fibrotic marker α-SMA were markedly decreased in *Kdm4d*-deficient HSC. Therefore, our research revealed that KDM4D was indeed indispensable for HSC activation and liver fibrogenesis in a TLR4/MyD88/NF-κB-dependent manner. Thus, targeting KDM4D provides an alternative approach against HSC activation, further, hepatic fibrogenesis.

We report that KDM4D might exert a pro-fibrotic role, which opens new horizons for hepatic fibrosis interference. It is conceivable that the presence of KDM4D drives fibrotic signaling through induction of TLR4. Certainly, the underlying mechanisms by which KDM4D facilitates liver fibrogenesis are much more complicated than we studied here. Therefore, we cannot fully exclude other signaling pathways modulated by KDM4D in liver fibrosis and a ChIP-sequencing study is warranted to uncover more potential targets of KDM4D.

In conclusion, our current findings provide critical insights into the molecular mechanisms for the regulation of liver fibrosis in an epigenetic fashion. Our data highlight a previously unknown role for KDM4D, which shed light on how histone modification enzymes coordinate with transcription factors to regulate TLR4 expression and liver fibrosis. Thus, blockade of KDM4D-TLR4 signaling in HSCs may serve as a novel approach for the epigenetic interruption of liver fibrosis. Currently, there is no specific inhibitor for KDM4D, hampering the direct blockade of KDM4D-TLR4 signaling from the very beginning. However, our findings provide rationale for the screening of small-molecule compounds that target KDM4D as an interventional strategy to reverse liver fibrosis. Therefore, roles of KDM4D in the context of liver fibrosis may merit thorough investigation in follow-up studies.

## Acknowledgements

We thank Asia-Vector Biotechnology Co. Ltd (Shanghai, China), Shanghai Tuoran Co. Ltd (Shanghai, China) and Shanghai Genechem Co. Ltd (Shanghai, China) for their technical support. We thank Dr. Michael Patrick for manuscript polishing and Dr. Linli Yang for technical assistance with HSC isolation.

## Funding

The research was supported by grants from Shanghai New Hundred Talents Program (No: XBR2013091), Shanghai Municipal Commission of Health and Family Planning, Key Developing Disciplines Program (No: 2015ZB0501), Shanghai Key disciplines program of Health and Family Planning (No: 2017ZZ02010) and Shanghai Sailing Program (No: 17YF1405200).

## Potential conflict of interest

Nothing to report.

## SUPPLEMENTARY MATERIAL ONLINE

Supplementary materials and methods

Figure S1. KDM4D knockdown suppresses HSC activation *in vitro*.

Figure S2. TLR4 expression is modulated by KDM4D in HSCs.

Figure S3. KDM4D regulates liver fibrosis through TLR4 signaling pathway.

Table S1: RT-qPCR primer sequences for mouse used in this study

Table S2: RT-qPCR primer sequences for LX2 (human) used in this study

Table S3: RT-qPCR primer sequences for T6 (rat) used in this study

Table S4: siRNA sequences targeting *KDM4D* in LX2 cells

Table S5: siRNA sequences targeting *Kdm4d* in mouse HSC

Table S6: siRNA sequences targeting *Kdm4d* in T6 cells

Table S7: siRNA sequences targeting *TLR4* in LX2 cells

Table S8: Heatmap data for Toll-like receptor signaling pathway

Table S9: Differentially expressed genes in KDM4D knockdown LX2 cells

Table S10: Antibodies used for western blotting in this study

Table S11: Antibodies used for ChIP in this study

